# Correlative evaluation of anticancer effects of a modified methioninase MGL-KPV using 2D and 3D cell models, human cancer xenografts in zebrafish embryos and Balb/c nude mice

**DOI:** 10.64898/2025.12.23.696173

**Authors:** Irina I. Khan, Saida Sh. Karshieva, Darina V. Sokolova, Chingis M. Aidossov, Yulia A. Borisova, Nikolay A. Bondarev, Anna A. Kudryavtseva, Sergei V. Bazhenov, Ilya V. Manukhov, Vadim S. Pokrovsky

## Abstract

**Purpose:** The emergence of new models of tumor growth requires a comparison of the results of the evaluation of the anticancer effect of single drug candidate for the selection of a relevant model.

**Methods:** We assessed the anticancer activity of the modified enzyme MGL-KPV using four models: 2D cultures, 3D spheroids, zebrafish embryo xenografts, and Balb/c nude mice. Cytotoxicity was tested in cancer cell lines and fibroblasts; efficacy was evaluated in zebrafish and mice. The effect of MGL does not require direct contact with cancer cells, making it unique drug candidate for *in vitro* and *in vivo* comparison.

**Results:** MGL-KPV showed selective cytotoxicity with IC_50_ values of 0.23–2.21 U/mL (2D) and 1.43–4.82 U/mL (3D), sparing fibroblasts. HCT-116 and Panc-1 were most sensitive. In zebrafish xenografts, tumor size reduced by 70% (HCT-116) and 43% (Panc-1); in mice — by 45% and 34%, respectively.

**Conclusions:** MGL-KPV demonstrated significant anticancer activity across all models. These results support step-by-step non-clinical proof-of-concept studies approach for amino-acid cleaving enzymes with anticancer properties.

## 1 INTRODUCTION

Modern non- and pre-clinical studies of anticancer compounds employ a wide range of non-clinical models — from conventional 2D cell cultures to advanced 3D systems and multiple *in vivo* platforms — each offering unique properties but also presenting specific limitations. 2D monolayer cultures remain nonredundant for high-throughput screening due to their simplicity and reproducibility, yet they fail in simulating key tumor microenvironment interactions like cell-cell contacts and extracellular matrix (ECM) architecture [1]. 3D *in vitro* models, such as spheroids, organoids, scaffold-based cultures, and microfluidic “organ-on-chip” platforms, more closely replicate spatial organization, cellular heterogeneity, hypoxia, and ECM interactions [2]. Despite their benefits, 3D models vary in complexity, standardization, and scalability, emphasizing the need for systematic cross-model comparisons to clarify their translational value [3].

Among *in vivo* models, human cell-derived xenografts in immunodeficient mice remain the gold standard. Zebrafish embryo models are gaining popularity, driven by 3Rs principles and the need for more ethical, scalable, and affordable approaches [4]. Their rapid development, optical transparency, and naturally immunodeficient early stage make them ideal for engrafting human tumor cells without immunosuppression.

This model allows real-time tracking of tumor growth, angiogenesis, and metastasis, and fits well into high-throughput drug screening pipelines [5]. Both zebrafish and mouse xenografts are now recognized in regulatory guidelines (FDA, EMA), with emphasis on using multiple models for better translational relevance. Cross-validation across 2D, 3D, and *in vivo* platforms helps uncover context-dependent drug responses and improves model selection in early-stage drug development.

In this context, our prior studies have characterized aminoacide-cleaving enzymes as anticancer agents, including asparaginase, L-lysine-α-oxidase, and methionine γ-lyase (MGL) [6– 8]. Here, we focus on MGL-KPV, a modified MGL variant bearing the KPV tripeptide derived from α-MSH, previously reported to confer anti-inflammatory activity [9] and mitigate hepatotoxicity. This work evaluates MGL-KPV’s cytotoxicity and anticancer efficacy across 2D and 3D *in vitro* models, and *in vivo* in human tumor xenografts in zebrafish embryos and Balb/c nude mice.

## 2. RESULTS AND DISCUSSION

### 2.1 Cytotoxic effect of MGL-KPV on 2D cancer cells cultures

To test MGL-KPV’s cytotoxicity *in vitro*, we used HCT-116, Huh7, Panc-1, LNCaP, and MCF-7 cancer cell lines. Past studies show gastrointestinal cancer cell lines are sensitive to KPV [9]. Normal human fibroblasts were included to evaluate selectivity.

Fibroblasts were the least sensitive to MGL-KPV, with IC_50_ of 2.2 U/mL in 2D and 9.1 U/mL in 3D cultures. Panc-1 and HCT-116 cells showed the strongest cytotoxic effects in 2D.

MGL-KPV was toxic to all cancer lines, with IC_50_ values between 0.23 and 2.21 U/mL (Table 1).

**Table 1.** Comparative table of the indicators of anticancer activity of MGL-KPV *in vitro* and *in vivo*

|  | *In vitro* cell models (IC<sub>50</sub>) |  | *In vivo* xenograft models (TGI<sub>max</sub>) |  |
| --- | --- | --- | --- | --- |
|  | 2D monolayer | 3D spheroid | Zebrafish embryo | Balb/c nude mice |
| HCT-116 | **0.83±0.08** | **1.43±0.28** | 70% | 45% |
| Panc-1 | **0.23±0.03** | **1.50±0.14** | 43% | 34% |
| Huh-7 | 1.93±0.16 | 3.92±0.30 | ND | ND |
| MCF-7 | 0.65±0.07 | 2.77±0.26 | ND | ND |
| LNCaP | 0.75±0.11 | 4.82±0.35 | ND | ND |
| Human normal fibroblast | 2.21±0.10 | 9.10±0.63 | ND | ND |

### 2.2 Antiproliferative effect of MGL-KPV on 3D spheroids

The lowest IC_50_ values were in HCT-116 (1.43 µM) and Panc-1 (1.50 µM) spheroids; Huh7 (3.92 µM) and LNCaP (4.82 µM) showed moderate sensitivity. Normal fibroblast spheroids had a much higher IC_50_ (9.1 µM), confirming MGL-KPV’s selective toxicity, consistent with 2D data (Table 1).

### 2.3 Assessment of MGL-KPV toxicity in *Zebrafish* embryos

Toxic effects of MGL-KPV appeared within 24 hours in *Danio rerio* embryos. At 10 U/mL, embryos showed delayed development, spinal deformities, and yolk sac swelling, lasting through the next day; all embryos died by 72 hours. At 100 U/mL, developmental delays and pericardial hemorrhages led to death by 48 hours. The LC50 was 3.2±1.1 U/mL. A non-toxic dose of 1.6 U/mL was chosen for therapeutic testing.

### 2.4 Anticancer effect of MGL-KPV in the HCT-116 and Panc-1 xenograft models using Zebrafish embryos

Cytotoxicity testing in 2D and 3D cultures identified HCT-116 and Panc-1 as the most sensitive lines to MGL-KPV, so they were selected for zebrafish xenografts.

MGL-KPV treatment significantly reduced HCT-116 tumor size in zebrafish embryos. After 24 hours, tumor area shrank 2.7-fold (0.032 ± 0.012 mm^2^ vs 0.086 ± 0.026 mm^2^ in controls; *p* < 0.001; Figure 1a, TGI = 63%). At 48 hours, the reduction was 3.3-fold (0.047 ± 0.063 mm^2^ vs 0.135 ± 0.029 mm^2^; *p* < 0.001; Figure 1b, TGI = 70%).

**Figure 1.**
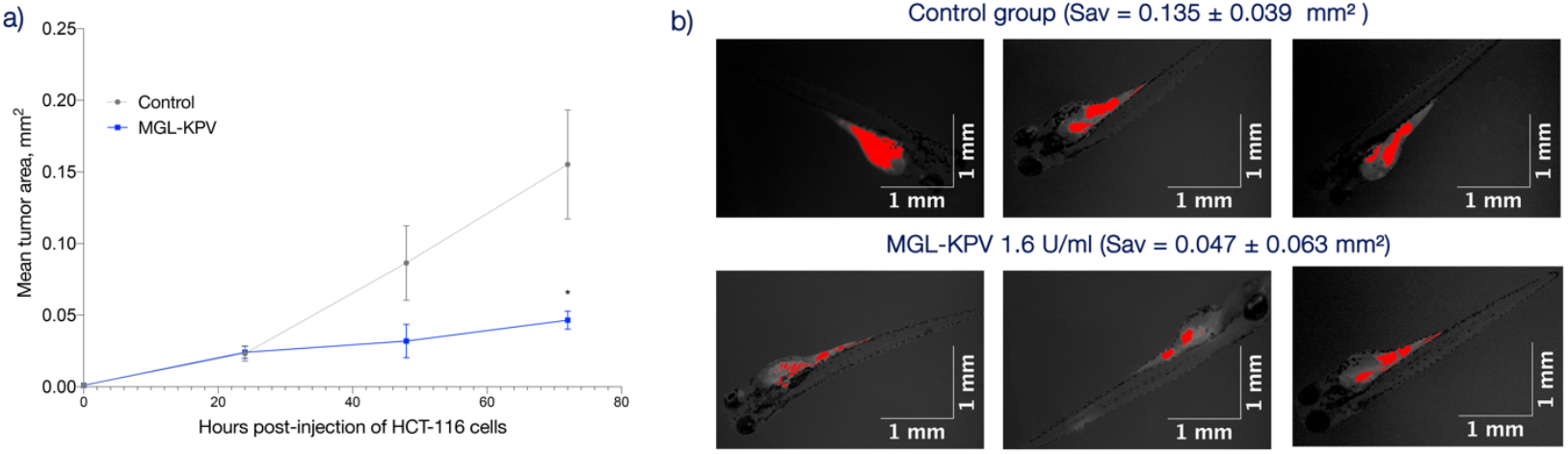
**(a)** Growth dynamics of HCT-116 xenografts in zebrafish embryos (*p < 0.001). **(b)** Images of zebrafish embryos with implanted human HCT-116 tumor cells (red) 24 hours after MGL-KPV treatment, compared to controls (showing smallest, largest, and average tumors). After 48 hours, tumors in treated embryos were 3.3 times smaller than controls. Average tumor area was 0.047 ± 0.063 mm^2^ vs. 0.135 ± 0.039 mm^2^ (p < 0.001). Images were processed by splitting color channels and applying a green filter to highlight tumors

Although MGL-KPV treatment resulted in a reduction in tumor burden in the Panc-1 zebrafish xenografts, the difference did not reach statistical significance (Figure 2a). The anticancer effect was assessed at 72 h post injection (hpi), with a TGI of 43%, and a mean tumor area of 0.031 ± 0.09 mm^2^ compared to 0.054 ± 0.016 mm^2^ in the control group (p=0.329; Figure 2b).

**Figure 2.**
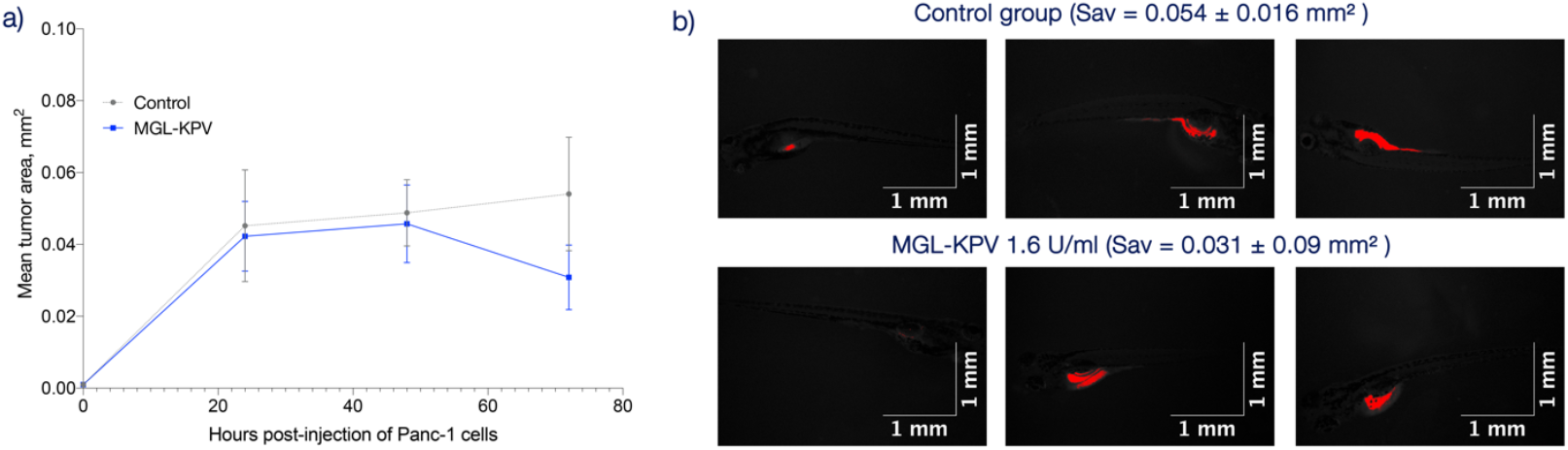
**(a)** Growth dynamics of Panc-1 xenografts in zebrafish embryos. **(b)** Images of zebrafish embryos with implanted human Panc-1 tumor cells (red) 24 hours after MGL-KPV treatment, compared to controls (smallest, largest, and average tumors). After 48 h, tumors in treated embryos were about 3.3 times smaller than controls. Average tumor area was 0.031 ± 0.09 mm^2^ vs 0.054 ± 0.016 mm^2^ in controls (p = 0.329). Images were processed by splitting color channels and applying a green filter to highlight tumors

### 2.5 Safety in Balb/c mice

MGL-KPV at studied doses (150, 300, or 600 U/kg intraperitoneally or 250, 500, or 1000 intravenously) did not exhibit any toxic effects nor did it cause observable behavioral changes or significant body weight loss in mice within a week post-treatment.

### 2.6 Anticancer effect of MGL-KPV in HCT-116 and Panc-1 Balb/c nude xenografts

Balb/c nude mice with HCT-116 xenografts started treatment on day 9 after cell implantation. Both doses (75 and 150 U/kg) showed dose-dependent tumor growth inhibition (TGI), peaking at 42% (p=0.093) and 45% (p=0.386) by day 3 post-treatment, with tumor volumes of 408.3 ± 77.7 mm^3^ and 387.6 ± 111.0 mm^3^ vs. 700.3 ± 157.8 mm^3^ in controls (Figure 3a).

**Figure 3.**
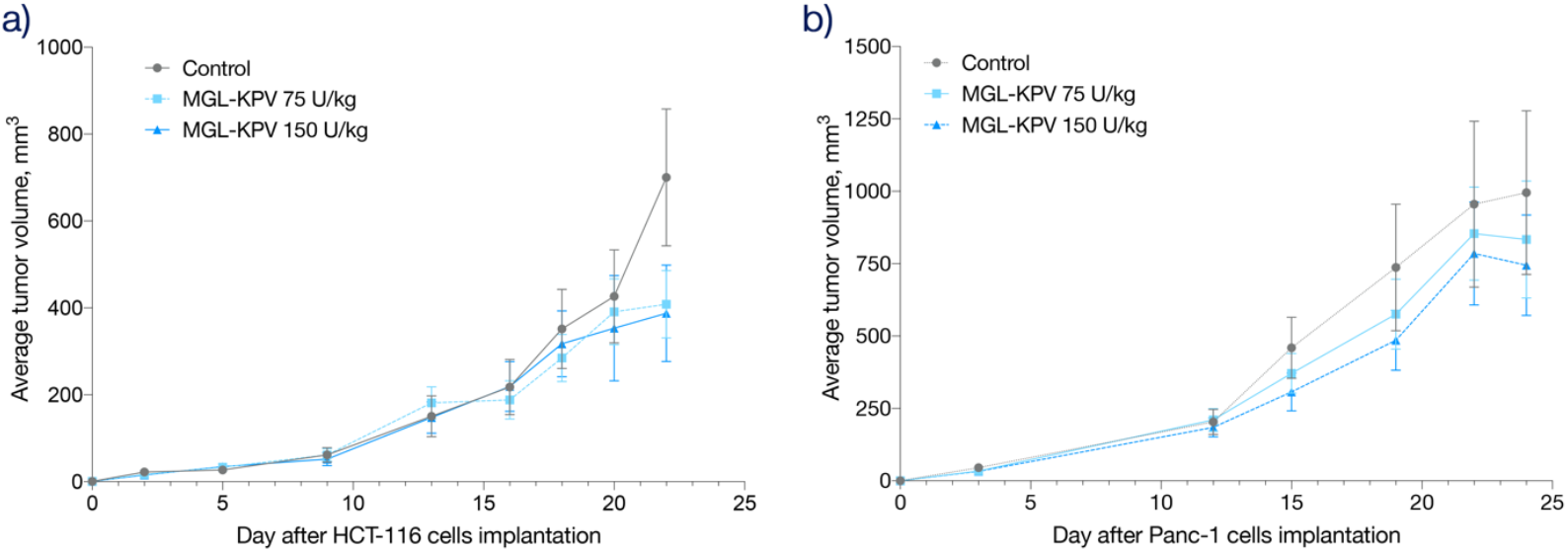
The dynamics of HCT-116 (a) and Panc-1 (b) xenograft growth in Balb/c nude mice treated with MGL-KPV 75 or 150 U/kg

Mice with Panc-1 xenografts began treatment on day 13. Tumor growth inhibition peaked on day 6, reaching 22% (p=0.790) at 75 U/kg and 34% (p=0.401) at 150 U/kg, with tumor volumes of 575.5 ± 120.8 mm^3^ and 485.7 ± 103.9 mm^3^, compared to 736.8 ± 218.7 mm^3^ in controls (Figure 3b).

## 3 DISCUSSION AND CONCLUSION

Methionine dependence is common in most solid cancers, making methionine-elimination therapy a promising anticancer approach, similar to L-asparaginase use in leukemia [10]. MGL (METase, EC 4.4.1.11) is a pyridoxal phosphate-dependent enzyme that catalyzes γ-elimination of L-methionine and β-elimination of L-cysteine and its derivatives. It also performs γ- and β-substitutions on methionine. MGL from *C. freundii* [11] and *Cl. sporogenes* [8] was reported to have a strong anticancer potential. It’s been used both monotherapy and in combination therapies [8, 12], including enzyme-prodrug approaches with sulfoxides [13]. The anticancer effect of MGL does not require direct contact with cancer cells, making it unique drug candidate for *in vitro* and *in vivo* comparison. MGL-KPV is a modified version of the native enzyme, designed to improve safety while retaining methionine-depleting activity. The KPV fragment may enhance anticancer effects by engaging the PepT1 transporter and modulating AKT/MAPK/NF-κB signaling [14].

KPV also inhibits TNF-α, likely by suppressing NF-κB activation [15, 16], which leads to DNA fragmentation in MCF-7 cells [17]. In Panc-1 cells, NF-κB regulates apoptosis [18], and targeting this pathway in HCT-116 cells has shown therapeutic promising potential [19].

In this study, we used four models: 2D cultures, 3D spheroids, zebrafish embryo xenografts, and Balb/c nude mouse xenografts (Table 1) to evaluate MGL-KPV’s anticancer activity.

In 2D cell models, MGL-KPV demonstrated cytotoxicity against all tested cancer lines at 0.2–2 U/mL. This matches results for MGL derived from *Aspergillus fumigatus*, which showed an IC_50_ of 1.452 U/mL in HCT-116 cells [20]. Similar active ranges were reported for native and mutant MGL from *C. freundii* and *C. novyi* [21].

Huh-7 cells were expected to be the most responsive to MGL-KPV due to high PepT1 expression [22]. However, notable effects were also seen in PepT1-negative lines, including HCT-116, Panc-1, Huh-7, MCF-7, and LNCaP. This suggests MGL-KPV acts through multiple mechanisms: methionine depletion, PepT1-mediated transport, and likely NF-κB inhibition via KPV.

Spheroids more accurately replicate key features of solid tumors-like hypoxia and ECM barriers — which can affect MGL-KPV activity [23]. In our study, cell viability in treated spheroids didn’t correlate with their size or growth. Sensitivity to MGL-KPV decreased in 3D models compared to 2D, with 1.7-to 6.5-fold reduction of the efficacy (Table 3). The spheroid model offered a more realistic platform for assessing anticancer activity, avoiding the overestimation of the effect often seen in 2D cultures.

HCT-116 and Panc-1 were the most responsive lines, consistent with prior data on C115H-MGL in Panc-1 xenografts [24].

In zebrafish embryos, MGL-KPV produced a stronger antitumor effect against HCT-116 xenografts (TGI_max_ = 70%) than in mice (TGI_max_ = 45%). Tumor suppression in zebrafish occurred faster and at lower doses, underscoring their value as a sensitive, efficient *in vivo* model for early drug screening.

Against Panc-1 xenografts, MGL-KPV showed modest effects in both systems. Still, the zebrafish model reached a TGI_max_ of 43%, slightly above the 34% observed in mice. The earlier onset and stronger effect in zebrafish may stem from better bioavailability or microenvironmental differences that enhance drug response.

Our temporary assumption is that anticancer activity *in vivo* correlated with *in vitro* results: effect in HCT-116 xenograft models were higher than in Panc-1. As the experimental models increased in complexity — from spheroids to xenografts in zebrafish embryos and Balb/c nude mice the efficacy of MGL-KPV gradually decreased. This likely relates to the tumor microenvironment (TME), with its complex cell interactions, variable pH, and enzyme inhibitors reducing enzyme efficiency and bioavailability [25]. Our observations shows that while 2D cultures are good for initial screening, they may overestimate efficacy.

Taken together, these results confirm the anticancer potential of MGL-KPV, which is reported for the first time. Improving enzyme stability and targeted delivery will be crucial to translating this approach into clinical settings.

## 4. EXPERIMENTAL

### 4.1 MGL-KPV preparation

The megL gene from *Clostridium sporogenes* was amplified using primers KPV MGL 5’ - TTGCATTGAATTCTTACACACCGGCTTAACTTAACTATTAAATCTAAAGCTTGTTT - 3’ and MGL reverse 5’-GTTTGCCACACAAAGGCTATACATACATGGA-3’ using plasmid pET-MGL-Sporog as a matrix [26]. The resulting fragment and the pET-MGL-Sporog plasmid were processed with PstI and EcoRI restriction endonucleases and ligated. The resulting fragment was processed with PstI and EcoRI restrictases and ligated with the pET-MGL-Sporog plasmid restricted to the same sites. The resulting pET-MGL-KPV construct contains a gene encoding MGL with three additional amino acids KPV at the C-terminus, under the control of the T7 promoter. *E. coli* BL21 (DE3) cells transformed with pET-MGL-KPV were used for biosynthesis, isolation and purification of MGL-KPV enzyme with three additional KPV amino acids. Small scale biosynthesis of enzyme was performed in autoinduction medium in flask [27]. The ANCUM-2 fermentation system (working volume 10 L; Institute of Biological Instrumentation of RAS, Pushchino) was used to produce the MGL-KPV pilot batch [8, 28]. Cell biomass after fermentation was resuspended in 10 mM potassium phosphate buffer supplemented with 1 mM EDTA, 0.2 mM PMSF and 1 μM pyridoxal phosphate (buffer A), lysed with a microfluidizer and centrifuged at 15,000 g for 40 minutes. The resulting clarified lysate was heat-treated with alcohol at 35°C with alcohol added up to 20% and spun again at the same parameters. The resulting solution was then applied by batch method overnight to a Q sepharose ion exchange resin pre-equilibrated in buffer A. The next day, the column was washed sequentially with 2 CV (column volume) of buffer A supplemented with 0.5% Triton-X114, followed by 2 CV buffer A with addition of 5% isopropanol, and then just 2 CV buffer A. The target protein was then run off under a KCl gradient and the fractions were analyzed with SDS PAGE. In the next step, fractions with highest MGL-KPV content were subjected to a two-step purification using ammonium sulfate. In the first step, non-target proteins were precipitated by adding ammonium sulfate to 35% percent of the saturation. The target protein was then precipitated by adding ammonium sulfate to 70% percent of the saturation. The precipitate was diluted in PBS and subjected to gel filtration for buffer change on a pre-equilibrated PBS Hiprep 26/10 Desalting column. The resulting protein solution was sterilized by filtration through 0.22 μm filters. Lyophilization and storage of the preparation was carried out as described in [29]. Stock solutions of the preparation were prepared *ex tempore* by dilution in physiological saline to a concentration of 20.85 mg/mL. MGL-KPV stock solutions were stored at +2-8°C for no more than 7 days. The specific activity of MGL-KPV was 24 units per mg of protein.

### 4.2 Cell lines

Human cancer cell lines MCF7 (breast, ATCC HTB-22), LNCaP (prostate, ATCC CRL-1740), HCT-116 (colon, ATCC CCL-247), Huh7 (liver, JCRB0403), Panc-1 (pancreas, ATCC CRL-1469), and non-tumor human dermal fibroblasts (HDF, Lonza CC-2511) obtained from the N.N. Blokhin National Research Medical Center collection with short tandem repeat-based (STR) authentifications and mycoplasma-free were used. Cells were cultured in DMEM (Gibco, USA) with 2 mM L-glutamine (PanEco, Russia), antimycotic antibiotic (Gibco), and 10% fetal bovine serum (Gibco), at 37°C in 5% CO2.

### 4.3 Cultivation of spheroids

Spheroids were formed in 96-well ultra-low attachment plates (Corning, USA) by seeding 1,000 cells/well in 100 μl. Spheroid size and roundness were analyzed with ImageJ.

### 4.4 Assessment of cytotoxicity of MGL in 2D and 3D cell models

To assess MGL-KPV cytotoxicity, monolayer cells (5×10^3^/well) and spheroids (1×10^3^/well) were seeded in 96-well plates and incubated overnight at 37°C in 5% CO2. MGL-KPV (0.2–50 U/mL) was added after 24 h (monolayer) or 48 h (spheroids). After 72 h, viability was measured by resazurin assay: medium replaced with 0.01% resazurin in DMEM, incubated ≥2 h, then fluorescence recorded at 560/590 nm.

### 4.5 Cultivation of zebrafish

Adult AB zebrafish were maintained at 28°C in aquaria with a 14-hour light and 10-hour dark cycle. Embryos were raised in E3 medium at 28°C until they reached the desired embryonic stage. Any unfertilized eggs or embryos with significant developmental defects 24 h after fertilization were detected under a Nexcope NSZ-810 microscope (Ningbo Yongxin Optics Co., Ltd, China) and removed from the experiment.

### 4.6 Zebrafish Embryo Acute Toxicity Test

Studies were conducted on zebrafish in accordance with [30]. 24 hpf wild-type AB zebrafish embryos were dechorionized and placed in 24-well plates. MGL-KPV to the wells in concentrations ranging from 0.1–100 U/ml in each well, with each concentration tested in triplicate. Each well procedure was performed in triplicate (n = 5 per group).

The embryos were examined 24, 48 and 72 hours after the addition of MGL-KPV. Morphological changes, including malformed yolk sacs, tail malformations, delayed development and impaired motor activity, were observed. To estimate the full range of mortality, embryonic deaths were counted at 96 hpf.

### 4.7 Zebrafish xenograft model and xenograft exposure assay

For xenotransplantation, 48 hpf wild-type AB zebrafish embryos were dechorionated and anesthetized in 0.02 mg/mL tricaine (Sigma-Aldrich). Approximately 200 CFSE-labeled HCT-116 or Panc-1 cells were injected into the yolk sac of each embryo. Embryos with visible cell masses were incubated at 32°C. Individual embryos (n = 6 per group) were placed into separate wells of a 48-well plate for monitoring.

At 24 h post-injection, embryos with visible fluorescent tumor cells were selected for treatment and incubated with 1.6 U/mL MGL-KPV or left untreated (control) for 48 hours.

Fluorescence imaging was performed at 0, 24, and 48 hours after treatment start. Tumor areas were quantified using ImageJ (NIH, USA) based on fluorescence intensity. Images were processed by splitting color channels and applying a green filter to enhance tumor visualization. Anticancer efficacy was assessed by comparing tumor areas between treated and control groups at each time point.

### 4.8 Safety test on Balb/c mice

Female Balb/c mice (22–28 g, 8–10 weeks old) were obtained from the N.N. Blokhin National Medical Research Center of Oncology. They were housed under controlled conditions (22 ± 2°C, 65 ± 5% humidity) with free access to food and water.

MGL-KPV was administered in a single dose of 150, 300, or 600 U/kg intraperitoneally, or 250, 500, or 1000 units per injection intravenously. Control mice received equal volumes of saline on the same schedule. Body weight was monitored daily for 7 days to assess toxicity and overall health.

### 4.9 Human xenograft tumor models

Balb/c nude mice were subcutaneously injected with 5×10^6^ HCT-116 or Panc-1 cells on both flanks. Tumor growth was monitored 2–3 times per week, with measurements every three days. Tumor volume (V) was calculated using length (L), width (W), and height (H).

Once average tumor volume reached 60 mm^3^ (HCT-116) or 180 mm^3^ (Panc-1), mice were randomized into three groups (n=6) and treated with MGL-KPV at 75 or 150 U/kg intravenously, daily for 10 days. Control mice received 0.9% saline on the same schedule.

The anticancer effect was evaluated by tumor growth inhibition (TGI).

### 4.10 Ethical statement

All animal experiments were conducted in accordance with the internationally accepted principles for laboratory animal use and care, as described in the EEC Directive of 1986; 86/609/EEC, and with approval from the Ethics Committee for Animal Experimentation of N.N. Blokhin Cancer Research Center (#2025-1b from 18th of February 2025).

*In vivo* data were analyzed in SPSS 21 (licensed June 26, 2013), with results shown as mean ± SE. Group comparisons used the Mann–Whitney test. Graphs were prepared in GraphPad Prism. Differences with p < 0.05 were considered statistically significant.

## ACKNOWLEDGEMENTS

MGL-KPV biosynthesis, purification, and lyophilization were performed with support of the Ministry of Science and Higher Education of the Russian Federation (agreement 075-03-2025-662, project FSMG-2025-0003). All other parts of the study were supported by the state program of the Ministry of Science and Higher Education of the Russian Federation (#075-01551-23-00; FSSF-2023-0006).

